# On tumoural growth and treatment under cellular dedifferentiation

**DOI:** 10.1101/2022.05.10.491343

**Authors:** Matthias M. Fischer, Nils Blüthgen

**Affiliations:** Institute for Theoretical Biology, Humboldt Universität zu Berlin, 10115 Berlin, Germany; Charité Universitätsmedizin Berlin, Institut für Pathologie, 10117 Berlin, Germany

**Author notes:** Corresponding author –.

**Keywords:** Cancer stem cell, Cellular hierarchy, Developmental trajectory, Hierarchical tissue, Non-stem cancer cell, Targeted therapy

## Abstract

Differentiated cancer cells may regain stem cell characteristics; however, the effects of such a cellular dedifferentiation on tumoural growth and treatment are currently understudied. Thus, we here extend a mathematical model of cancer stem cell (CSC) driven tumour growth to also include dedifferentiation. We show that dedifferentiation increases the likelihood of tumorigenenis and the speed of tumoural growth, both modulated by the proliferative potential of the non-stem cancer cells (NSCCs). We demonstrate that dedifferentiation also may lead to treatment evasion, especially when a treatment solely targets CSCs. Conversely, targeting both CSCs and NSCCs in parallel is shown to be more robust to dedifferentiation. Despite dedifferentiation, perturbing CSC-related parameters continues to exert the largest relative effect on tumoural growth; however, we show the existence of synergies between specific CSC- and NSCC-directed treatments which cause superadditive reductions of tumoural growth. Overall, our study demonstrates various effects of dedifferentiation on growth and treatment of tumoural lesions, and we anticipate our results to be helpful in guiding future molecular and clinical research on limiting tumoural growth in vivo.

## I. INTRODUCTION

Tumoural cell populations can consist of different subpopulations [1, 2]. Especially of note are cancer stem cells (CSCs), which are defined by their capacity for indefinite self-renewal and their ability to initiate clonal outgrowth [3]. In contrast, non-stem cancer cells (NSCCs) often show a limited proliferative capacity [4]. The discovery of this intratumoural heterogeneity has given rise to a promising paradigm of cancer treatment, in which therapy is specifically directed at eradicating CSCs, instead of indiscriminately targeting the complete tumour mass [5, 6].

Mathematical modelling has been used extensively to study the dynamics of tumoural growth; for a review refer to Yin *et al*. [7]. Multiple model structures which specifically integrate intratumoural heterogeneity have been proposed; in particular, authors have distinguished between proliferative and quiescent cells [8–10], or treatment-sensitive and -resistant cells [11–15]. Finally, Weekes *et al*. [16] proposed a multicompartment model consisting of indefinitely cycling CSCs and NSCCs with limited proliferative potential. This model has successfully been used by Werner *et al*. [17] to estimate the fraction of CSCs in growing tumours from patient-specific treatment trajectories.

Both healthy and cancer cell populations show a reversal of their developmental trajectories in which differentiated cells regain stem cell characteristics [18–25]. The effects of such a dedifferentiation on the homoeostasis of healthy tissue [26, 27], on the speed of mutation acquisition during carcinogenesis [28], on the phenotypic equilibrium [29, 30], on transient overshoots [31, 32], on the invasion of differentiated malignant cells into cellular hierarchies and on tumour survival in the limit of a small number of cells [34] have already been studied mathematically. While [35] have added a uniform cellular dedifferentiation rate for all non-stem compartments to the model by [16], they specifically fitted the model to a set of *in vitro* measurements on a colon carcinoma cell line and used the model to study the population dynamics after radiation treatment. An exhaustive mathematical analysis of the effects of dedifferentiation on growth and treatment of an arbitrary hierarchical tumour, however, has to the best of our knowledge not yet been performed.

Here, we aim to close this gap in the literature and elucidate how dedifferentiation affects the dynamics of hierarchically organised tumours. To this end, we extend the model of a cancer stem cell-driven tumour by Weekes *et al*. [16] to include dedifferentiation of arbitrary subsets of non-stem compartments, and study its properties both analytically and numerically. Importantly, we also use the model to exhaustively examine how dedifferentiation influences possible ways of causing tumour shrinkage.

## II. MATERIALS AND METHODS

### A. A model of tumoural growth with dedifferentiation

Our model is based on the work of Weekes *et al*. [16] and [35], whose notation we follow. We distinguish between a cancer stem cell (CSC) compartment *C*, and *M* non-stem cancer cell (NSCC) compartments *N*_1_, …, *N*_*M*_ with differing remaining proliferative capacity. CSCs divide indefinitely at a rate *K*_*C*_, and every division generates either two new CSCs, one CSC and one NSCC *N*_1_, or two NSCCs *N*_1_, with probabilities *P*_*C*_, *P*_*A*_, *P*_*D*_, respectively. In contrast, NSCCs only have a limited capacity of *M −* 1 divisions: NSCCs *N*_*i*_ divide at rate *K*_*N*_, thereby giving rise to two daughter cells *N*_*i*+1_ for *i* = 1…*M −* 1, and, in contrast to Weekes *et al*. [16], we assume that *N*_*M*_ cells are cell-cycle arrested. CSCs and NSCCs undergo apoptosis at rates *D*_*C*_, *D*_*N*_, respectively.

We add to this model the possibility of NSCCs *N*_1_…*N*_*d*_ dedifferentiating back into CSCs at a rate *K*_*T*_, where *d* can be any integer between 1 and *M*. Thus, our model is given by the following differential equations:

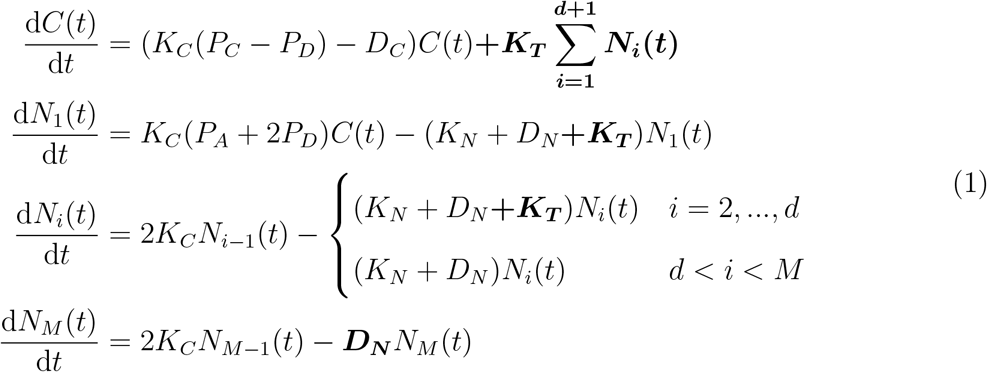

For better readability, we have set our changes to the original model by Weekes *et al*. [16] in boldface. A schematic overview of the extended model is presented in Figure 1A. The model suggests the following aggregate parameters which we will use throughout this work: The realised growth rate of the CSC compartment *β* = *K*_*C*_(*P*_*C*_ *− P*_*D*_) *− D*_*C*_, the flux *φ* = *K*_*C*_(*P*_*A*_ + 2*P*_*D*_) of cells from the CSC compartment into the *N*_1_ compartment, the rate *ε* = *K*_*N*_ + *D*_*N*_ at which cells exit the NSCC compartments *N*_1_…*N*_*M−*1_, and *δ* = 2*K*_*N*_, the rate at which cells enter NSCC compartments *N*_2_…*N*_*M*_ as a consequence of cell division in the respective preceding compartment.

**FIG. 1:**
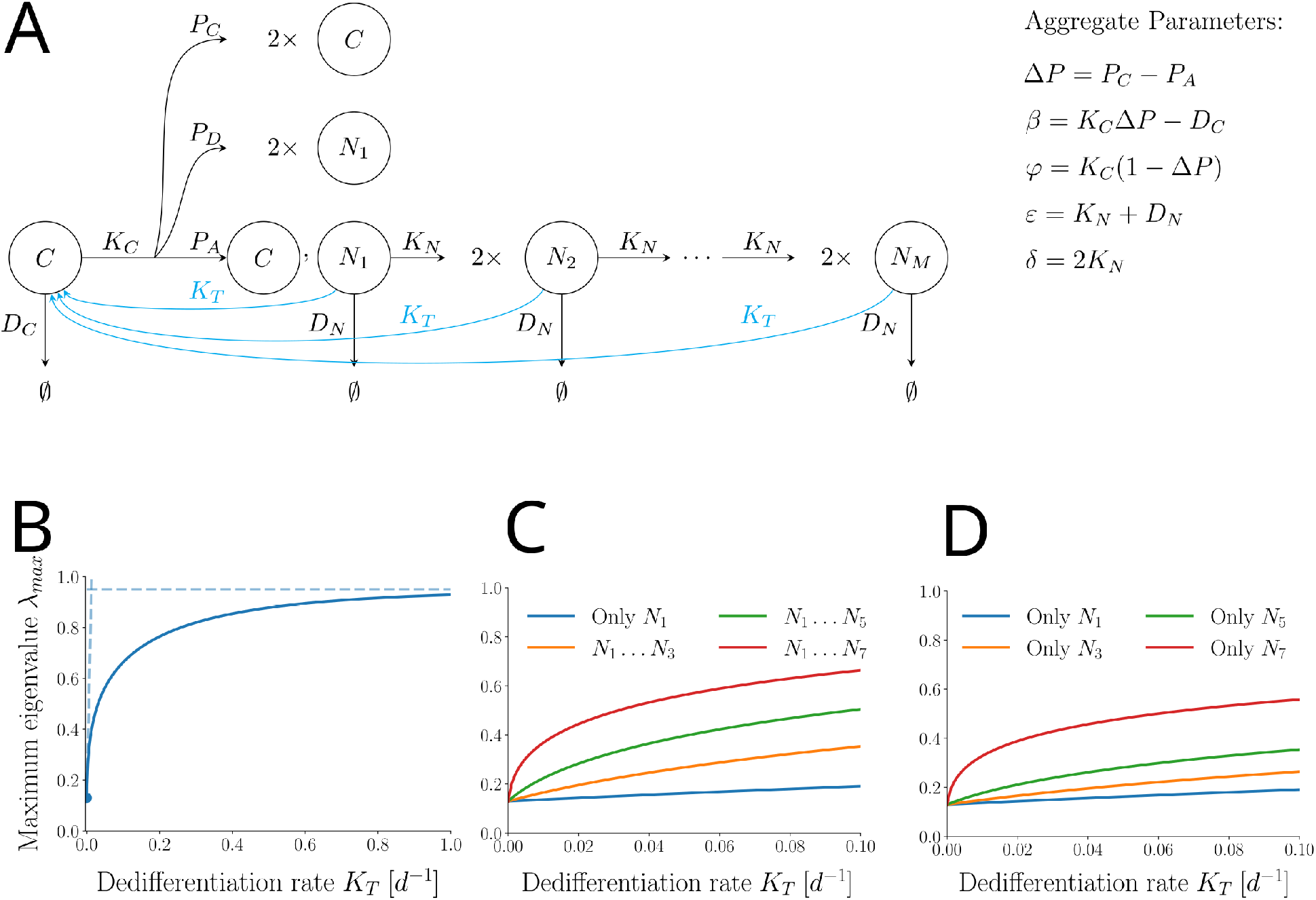
Our model of cancer stem cell-driven tumoural growth with dedifferentiation and an illustration of some of its analytically shown properties. **A** Schematic model sketch, with dedifferentiation marked in blue. **B** Dedifferentiation increases the maximum eigenvalue of the system. Dashed lines are asymptotics for small and large dedifferentiation rates *K*_*T*_ based on analytical estimates. **C** If more NSCC compartments dedifferentiate, this effect becomes stronger. **D** Later NSCC compartments contribute more strongly to the increase in maximum eigenvalue.

In particular, throughout this work we will consider two special cases of this general model. First, the special case of *d* = 1, i.e. a tumour in which only the youngest NSCCs *N*_1_ are able to dedifferentiate back to a stem-like cell state. This case is analytically easy to treat and may be a reasonable biological assumption. Second, we will also work with the case of *d* = *M*, i.e. a tumour in which all NSCC compartments dedifferentiate. An analytic treatment of this model structure is possible, however it yields expressions too complicated for a meaningful interpretation. For this reason, we will treat this case by using analytical matrix perturbation theory to approximate the effects of sufficiently small values of *K*_*T*_ (see Section III).

### B. Standard parametrisation of the model

We follow the standard parametrisation provided in Weekes *et al*. [16], which we summarise in Table I. The order of magnitude of the NSCC dedifferentiation rate *K*_*T*_ can be gauged from Wang *et al*. [35] who determined *K*_*T*_ *≈* 0.269*d*^*−*1^ in the human colon carcinoma cell line SW620 *in vitro*. However, under standard parametrisation this value would lead to a biologically implausible rate of overall tumour growth of 0.805 *d*^*−*1^. For this reason, we limited *K*_*T*_ to the interval of [0; 0.1] *d*^*−*1^, with a standard value of 0.05 *d*^*−*1^.

**TABLE I:** The standard parametrisation of the model from Weekes *et al*. [16]. CSC: cancer stem cell. NSCC: non-stem cancer cell.

| Symbol | Description | Value | Range | Unit | Comment |
| --- | --- | --- | --- | --- | --- |
| **Cell-cycling rates** |  |  |  |  |  |
| $K_C$ | CSC cycling rate | 1.0 | 0.5 – 1.5 | $d^{-1}$ | |
| $K_N$ | NSCC cycling rate | 1.0 | 0.5 – 1.5 | $d^{-1}$ | |
| **CSC fate probabilities** |  |  |  |  |  |
| $P_S$ | Probability CSC doubling | 0.18 | 0.01 – 0.35 | num. | |
| $P_A$ | Probability asymm. CSC division | $1 - P_S$ | – | num. | |
| $P_D$ | Probability double CSC differentiation | 0 | – | num | |
| **Apoptosis rates** |  |  |  |  |  |
| $D_C$ | CSC apoptosis rate | 0.05 | 0 – 0.1 | $d^{-1}$ | |
| $D_N$ | NSCC apoptosis rate | 0.05 | 0 – 0.1 | $d^{-1}$ | |
| **Other** |  |  |  |  |  |
| $M$ | Nb. NSCC divisions | 7 | 3 – 11 | num. | |
| $K_T$ | NSCC dedifferentiation rate | 0.05 | 0 – 0.1 | $d^{-1}$ | Systematically varied |

### C. Numerical examinations

#### 1. Virtual cohort trials

Virtual cohort patients were generated by uniformly drawing system parameters from the range of possible parameter values provided in Weekes *et al*. [16]. Thus, *D*_*C*_, *D*_*N*_ ∼𝒰 (0, 0.1), *M* ∼ 𝒰 (3, 11). For cell cycling ranges, only point estimates *K*_*C*_ = *K*_*N*_ = 1.0 *d*^−1^ were given, so we sampled from biologically plausible ranges as follows: *K*_*C*_, *K*_*N*_ ∼ 𝒰 (0.5, 1.5). For the CSC fate probabilities, using the constraint of *P*_*C*_ + *P*_*A*_ + *P*_*D*_ = 1, we introduced the aggregate parameter Δ*P* = *P*_*C*_ *− P*_*D*_, which we sampled Δ*P* ∼ 𝒰 (0.01, 0.35) following the parametrisation by Weekes *et al*. [16]. Dedifferentiation rate *K*_*T*_ was varied systematically in order to study its effect, or sampled *K*_*T*_ ∼ 𝒰 (0, 0.1).

We excluded a virtual patient if *λ*_*max*_ < 0 (negative tumoural growth) or *λ*_*max*_ > 0.2 (biologically implausibly fast tumoural growth). We also excluded those virtual patients, in which the CSC compartment had a negative growth (*K*_*C*_Δ*P* − *D*_*C*_ < 0), and thus the simulated tumour would only be able to grow due to a sufficient amount of dedifferentiation, which we deemed biologically unlikely.

We examined the effects of the following treatments, both isolated and in combination: First, causing CSC or NSCC apoptosis at a rate randomly sampled from *𝒰* (0, 1), modelling the effect of a cytotoxic treatment. Second, decreasing the CSC or NSCC cycling rate by a fraction sampled from *𝒰* (0, 1) times its initial values, modelling the effect of a cell-cycle inhibitor. Third, reducing the CSC fate parameter Δ*P* to a value sampled from *U* (*−*1, Δ*P*_0_), where Δ*P*_0_ represents the initial value of Δ*P*, to model the effect of a differentiation-inducing pharmaceutical.

#### 2. Single parameter sensitivities

To account for the different units and baseline values of the system parameters, we calculate the dimensionless, relative sensitivities (or “elasticities”) of the model parameters [36]. These quantify the ratio between an infinitesimal relative (i.e. fractional) change in one system parameter *X* and the corresponding relative change in the leading eigenvalue *λ*_*max*_ of the system (see Figure 5A for an illustration). Thus, we calculate

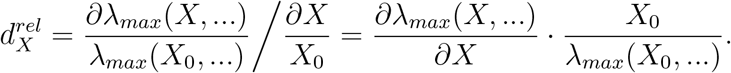

which we numerically approximate via the finite difference

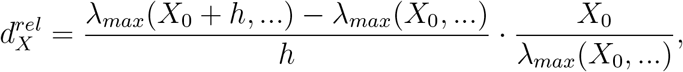

using *h* = 10^*−*6^.

#### 3. Treatment interactions

To numerically quantify the interaction of the alteration of two system parameters *X* and *Y*, Smith *et al*. [37] suggested the following formula based on second-order mixed partial derivatives:

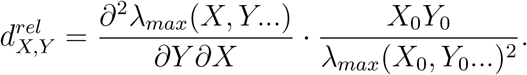

However, because the different treatments are applied into different directions (e.g. cell-cycling rates will be reduced, whereas apoptosis rates will be increased), the results of this calculation would not be straightforward to interpret intuitively: in some cases, a positive sign would hint at a synergistic effect, whereas for other treatment combinations the opposite would be true. For this reason, we slightly modify the previous approach and instead calculate the second derivative “into the direction of treatment” and approximate via the following finite difference scheme:

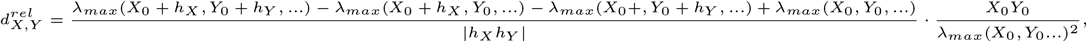

where *h*_*X*_ and *h*_*Y*_ are either 10^*−*6^ if the treatment associated with the respective system parameter increases the parameter (apoptosis rates *D*_*C*_, *D*_*N*_), or *−*10^*−*6^ if the parameter is decreased during treatment (cell-cycling rates *K*_*C*_, *K*_*N*_, CSC fate parameter Δ*P*, NSCC dedifferentiation rate *K*_*T*_). In this way, negative values always indicate a synergistic interaction, whereas positive values denote antagonistic (subadditive) treatment interactions.

#### 4. Implementation

All numerical examination have been implemented in the Python programming language [38], version 3.8. All computations have been carried out on a 64-bit personal computer with an Intel Core i5-3350P quad-core processor running Manjaro Linux, kernel version 5.10.56-1.

We provide the complete commented source code of all of our numerical examinations in the form of an iPython juypter notebook in the following github repository: https://github.com/Matthias-M-Fischer/Tumourgrowth.

## III. RESULTS

### A. Dedifferentiation alters tissue stability and tumour growth

We extended a literature model [16] of cancer stem cell-driven tumour growth to include dedifferentiation of non-stem cancer cells (NSCCs). A sketch of the model is provided in Figure 1A with dedifferentiation marked in blue. First, we study the influence of dedifferentiation on the existence and stability of the steady states of the model.

#### 1. Dedifferentiation may contribute to tumorigenesis and increases tumoural growth speed, both modulated by NSCC proliferative capacity

Regardless of our choice of *d* (the index of the last NSCC compartment that still dedifferentiates), the System 1 is obviously linear and homogeneous. Thus, we may write the system as

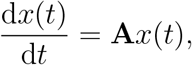

where *x*(*t*) = [*C*(*t*), *N*_1_(*t*), …*N*_*M*_ (*t*)]^*T*^ is the vector of compartment sizes, and **A** *∈* R*n×n* its constant coefficient matrix. Because **A** is nonsingular, the system permits exactly one equilibrium, which is the trivial steady state of tumour extinction given by 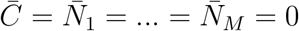. To asses the stability of the extinction state, we consider the eigenvalues of **A**.

The case of *d* = 1, where only the first NSCC compartment can dedifferentiate, allows for a direct calculation. Using the aggregate parameters introduced in Section II and shown in Figure 1A, we get:

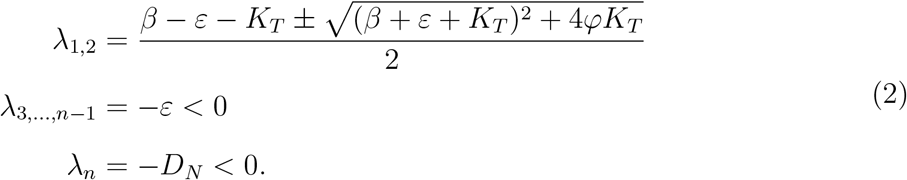

Clearly, if *K*_*T*_ = 0 we obtain a maximum eigenvalue of *λ*_*max*_ = *λ*_1_ = *β*. However, for every *K*_*T*_ *>* 0, due to *φ >* 0 the radicand will always exceed (*β* + *ε* + *K*_*T*_)^2^, causing the numerator to be strictly greater than 2*β*. Hence, *λ*_*max*_ *> β*. Thus, with dedifferentiation the maximum eigenvalue of the system will increase.

Next, we consider the model with arbitrary *d ≤ M*. In this case, a direct calculation of the eigenvalues of **A** for *K*_*T*_≠ 0 is possible, but leads to expressions too complicated for a direct analysis. For this reason, we use analytical matrix perturbation theory to approximate the leading eigenvalue for small values of *K*_*T*_ (see Appendix A). We find that the leading eigenvalue of the system follows a positive and increasing linear function of *K*_*T*_ :

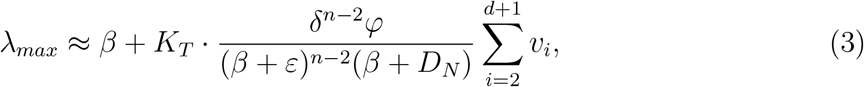

where *v*_*i*_ represents the *i*-th element of the leading eigenvector *v* of the system.

In this way, dedifferentiation can be responsible for *λ*_*max*_ exceeding zero, and thus for unbounded growth to occur in the first place. Additionally, if unbounded growth does occur, after a transient the system will grow at rate *λ*_*max*_. In this way, dedifferentiation will also increase tumoural growth speed.

Let 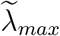 denote the increase in the leading eigenvalue *λ*_*max*_ compared to *β*, which arises as a consequence of dedifferentiation. Strikingly, under the assumption that the realised *in vivo* growth rate *β* of the CSC compartment and the NSCC apoptosis rate *D*_*N*_ are small compared to the NSCC cycling rate *K*_*N*_ (see Section II), we find that

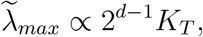

indicating that dedifferentiation causes a sharp linear increase of the leading eigenvalue with a slope that is exponential in the number of NSCC divisions (see the blue dashed line in Figure 1B for an illustration and Appendix A for details). Accordingly, limiting the number of NSCC divisions may be an interesting therapeutic approach for reducing tumoural growth.

This effect, however, will saturate for sufficiently large *K*_*T*_, as can be shown by using the Gershgorin circle theorem [39]: We can derive an upper bound of all eigenvalues of the system, finding that *∀i* : *λ*_*i*_ *≤* max*{β* + *φ, δ − ϵ}, i*.*e*. regardless of dedifferentiation rate and the number of dedifferentiating compartments, the growth of the system will be less or equal than the maximum of *{K*_*C*_ *− D*_*C*_, *K*_*N*_ *− D*_*N*_ *}* (see the blue horizontal dashed line in Figure 1B for an illustration). In other words, the growth-stimulating effect of dedifferentiation is bounded from above, which is biologically plausible as well.

#### 2. Different NSCC compartments contribute differently

In the previous section, we studied the effects of introducing dedifferentiation onto the system as a whole. Now we examine how the different NSCC compartments contribute to these effects. From Equation 3, one may easily see that increasing the number of dedifferentiating compartments will lead to more (strictly positive) summands being added to the leading eigenvalue, while all other terms stay constant. Thus, the more compartments are dedifferentiating, the greater the dedifferentiation-induced increase in *λ*_*max*_ will be.

As a numerical illustration of this finding, Figure 1C shows the leading eigenvalue *λ*_*max*_ of the system under standard parametrisation as a function of the dedifferentiation rate *K*_*T*_ for different subsets of NSCC compartments dedifferentiating. Note how a larger number of dedifferentiating NSCC compartments causes the leading eigenvalue to increase more strongly.

The ratios (*β* + *ε*)*/δ* and (*β* + *D*_*N*_)*/δ* can be expected to fall below unity (see Section II). Accordingly, the effect of later compartments dedifferentiating will be larger than the effect of earlier ones (see Equation 3). Figure 1D illustrates this finding under standard parametrisation, depicting the leading eigenvalue *λ*_*max*_ of the system as a function of the dedifferentiation rate *K*_*T*_, if only *N*_1_, *N*_3_, *N*_5_, or only *N*_7_ cells dedifferentiate. Note how every NSCC compartment exerts a stronger effect on the leading eigenvalue than its respective predecessors.

### B. Treatments with a single biological effect are more robust to dedifferentiation when CSCs and NSCCs are targeted in parallel

In the first part of this paper, we examined how the unperturbed system behaviour changes if non-stem cancer cells can dedifferentiate. Now, we turn to the question of potential ways to limit tumoural growth. To this end, we study the effect of parameter alterations on tumour growth rate and how the effects of such alterations are influenced by dedifferentiation. In this section, we focus on treatments which only exert one biological effect such as a reduced cell cycling rate, or an increased apoptosis rate.

#### 1. Treatment-resistant CSCs lead to therapy evasion

As a plausibility check, we start by analysing the effect of CSCs showing treatment resistance. In this case, a treatment can only alter NSCC parameters, but CSC parameters stay unaffected. Thus, in the absence of dedifferentiation (*K*_*T*_ = 0) the CSC compartment will grow according to *C*(*t*) = *C*(0)*e*^*βt*^ *>* 0. If additionally dedifferentiation takes place, the CSC compartment will grow at the same speed or faster due to the additional backflow of NSCCs into the CSC compartment (it is easy to prove that all *N*_*i*_(*t*) *≥* 0 by considering the system behaviour at the boundary). Hence, under dedifferentiation *∀t* : *C*(*t*) *≥ C*(0)*e*^*βt*^ *>* 0. Thus, as expected a resistance of CSCs to treatment causes continued growth of the tumour.

We also illustrate this numerically by studying a treatment which stimulates NSCC apoptosis *D*_*N*_ as an example (Figure 2A, left panel). No value for *D*_*N*_ will ever lead to long-term tumour extinction as shown above (illustrated in the middle panel). Integrating the system numerically with an exemplary treatment of *D*_*N*_ = 5 shows an initial reduction in tumour size, but the tumour will be replenished by surviving treatment-resistant CSCs (right panel). We also observe that during this regeneration the fraction of stem cells increases. These findings reproduce the clinical reality of e.g. chemotherapy first reducing tumour size, after which a relapse happens due to the survival and expansion of treatment-resistant CSC clones.

**FIG. 2:**
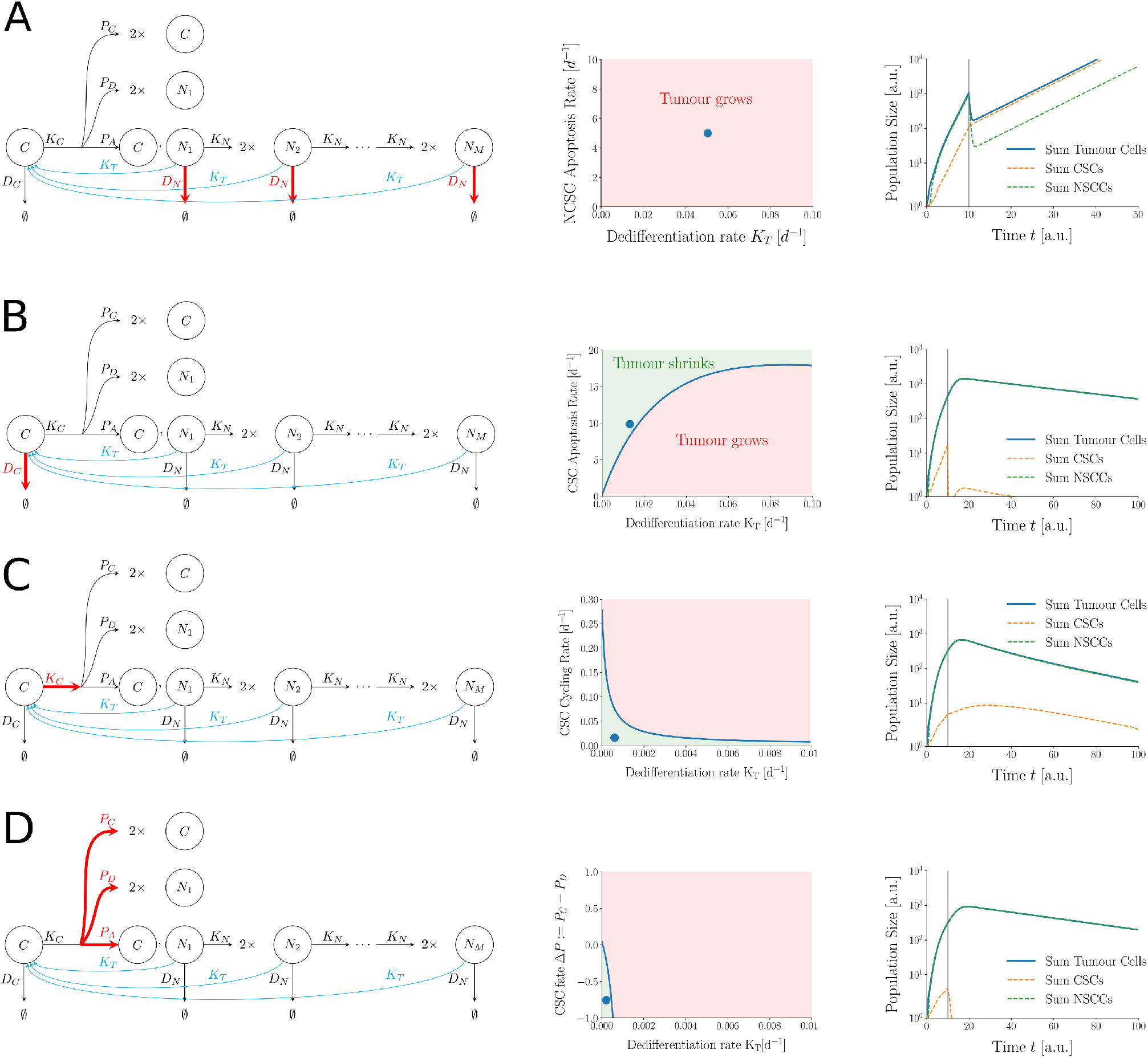
Controlling CSCs is necessary for treatment success; however, dedifferentiation may cause treatment failure. **A** Therapy-resistant CSCs cause treatment failure. Left: Illustration of exclusively targeting NSCC apoptosis rate *D*_*N*_ (marked in red) with unaffected CSC apoptosis rate *D*_*C*_ as a consequence of therapy-resistant CSCs. Middle: No value for *D*_*N*_ can lead to tumour extinction. Dot marks parametrisation for numerical simulation on the right. Right: Exemplary numerical simulation showing an initial reduction in tumour size as a consequence of NSCC apoptosis, and subsequent CSC-driven tumour relapse. Vertical grey line indicates onset of treatment. **B** Left: Targeting CSC apoptosis rate *D*_*C*_. Middle: Increasing dedifferentiation rate *K*_*T*_ requires a stronger apoptotic treatment for treatment success. Right: Exemplary numerical simulation of a successful treatment. **C** Left: Targeting CSC cycling rate *K*_*C*_. Middle: Dedifferentiation requires a stronger reduction of *K*_*C*_. Right: Exemplary numerical simulation. **D** Left: Targeting CSC fate probabilities. Middle: Dedifferentiation requires a stronger effect on CSC fate probabilities. Right: Exemplary numerical simulation.

#### 2. Targeting only CSCs can quickly fail in the face of dedifferentiation

Previously, we have confirmed that treatment-resistant CSCs cause treatment failure and tumour relapse, which is in accordance with clinical reality. Thus, we now examine the case, where CSCs are not resistant to treatment and we are thus able to target them. In this section, we only consider treatments that exclusively affect CSCs.

For the model where only *N*_1_ cells dedifferentiate, we find that only sufficiently strong treatments will be successful under dedifferentiation; however, for weaker treatments there always exists a critical value of *K*_*T*_ at which the treatment begins to fail. For instance, without dedifferentiation, introducing apoptosis in the CSC compartment succeeds as soon as *D*_*C*_ *> K*_*C*_Δ*P*. With dedifferentiation, however, we need at least *D*_*C*_ *> K*_*C*_ to guarantee treatment success in all cases. A weaker treatment may also succeed, but only if the dedifferentiation rate is sufficiently small:

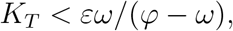

where *ω* := *D*_*C*_ *− K*_*C*_Δ*P* represents the strength of the apoptosis induction. For all calculations and all critical values, see Appendix B.

For the case of arbitrary *d*, we can again approximate the system behaviour, but only for sufficiently weak CSC-treatments. Those can all be evaded with a sufficiently large dedifferentiation rate *K*_*T*_. In contrast, for stronger treatments the approximation cannot be constructed because of the emergence of a multiple leading eigenvalue (see Appendix B).

Hence, we now examine numerically in more detail the exact susceptibility of CSC treatments to dedifferentiation. To this end, we use the standard parametrisation of the model (Table I) and systematically vary the dedifferentiation rate *K*_*T*_. In case of all three treatment avenues, we notice that even for small values of *K*_*T*_ stronger treatments become necessary to reach tumour shrinking (Figure 2B, C, D). Thus, a treatment regime which works in case of low *K*_*T*_ may fail for larger dedifferentiation rates. In this way, dedifferentiation can contribute to treatment evasion if the treatment is solely directed at the CSC compartment of the tumour.

#### 3. Targeting CSCs and NSCCs in parallel is more robust to dedifferentiation

In the previous section, we have observed that targeting only CSCs may easily fail in the face of dedifferentiation. Thus, we now examine whether targeting both CSCs and NSCCs in parallel can be more effective (see Appendic C for all details). We start with the model in which only *N*_1_ cells can dedifferentiate, finding that for all imaginable CSC treatments additionally inducing apoptosis in NSCCs can be expected to prove more robust to dedifferentiation. Interestingly, the same holds for increasing NSCC cycling rate *K*_*N*_, which might seem counterintuitive at first glance. However, this finding does make sense, because in the examined model only *N*_1_ cells dedifferentiate. Thus, increasing *K*_*N*_ leads to NSCCs leaving this compartment faster, and thus reduces cellular dedifferentiation.

For the case of an arbitrary number of dedifferentiating NSCC compartments, we can use the Gershgorin circle theorem [39] to derive exact upper bounds on all system eigenvalues for arbitrarily large *K*_*T*_ (see Appendix C for details). In this way, we find that a sufficiently strong treatment solely directed at CSCs can be guaranteed to succeed in the face of dedifferentiation if an additional, sufficiently strong treatment acts upon the NSCC compartments. If, however, only the CSC compartment is targeted, we cannot guarantee treatment success. In fact numerical examination carried out in the next section will demonstrate that in this case treatment evasion can happen easily even for small values of the dedifferentiation rate *K*_*T*_.

#### 4. Virtual cohort trials confirm robustness of synergistic effects of co-targeting CSCs and NSCCs

Now, we numerically examine in more detail the extent of these synergies for varying values of *K*_*T*_. We begin by using the standard parametrisation of the model. Targeting both CSC and NSCC apoptosis rates *D*_*C*_, *D*_*N*_ (sketched in Figure 3A) is indeed less sensitive to dedifferentiation as depicted in Figure 3B, where we plot those regions in parameter space, where treatment is successful. Note, how for small values of *D*_*N*_ an increase in dedifferentiation rate requires significantly higher CSC apoptosis rates *D*_*C*_ for a successful treatment; but this increase is diminished if in parallel the NSCC apoptosis rate is increased as well.

**FIG. 3:**
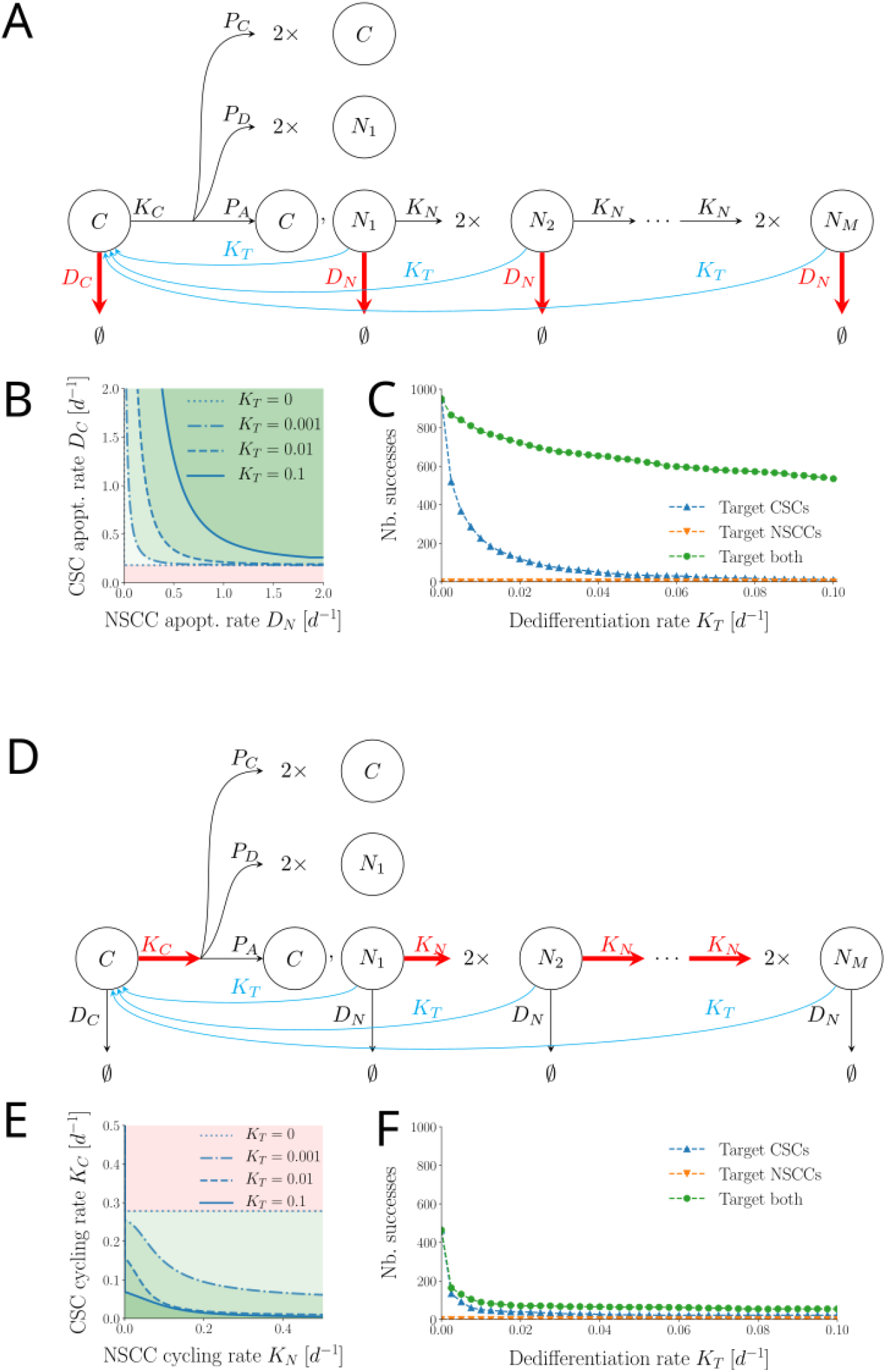
Targeting CSCs and NSCCs in parallel is more robust to dedifferentiation. Numerical results using standard parametrisation and a virtual patient cohort. **A** Illustration of a treatment stimulating both CSC apoptosis rate *D*_*C*_ and NSCC apoptosis rate *D*_*N*_. **B** Regions in (*D*_*C*_, *D*_*N*_) space under standard parametrisation where the tumour shrinks for different values of dedifferentiation rate *K*_*T*_. Note that a larger NSCC apoptosis rate *D*_*N*_ makes the treatment more robust to dedifferentiation. **C** Confirmation of the result in a virtual cohort trial with 1000 patients. Increasing both CSC and CSCC apoptosis rate leads to a higher fraction of cases with tumour shrinkage in the face of dedifferentiation. **D** Illustration of a treatment reducing both CSC cycling rate *K*_*C*_ and NSCC cycling rate *K*_*N*_. **E** Regions in (*K*_*C*_, *K*_*N*_) space under standard parametrisation where the tumour shrinks for different values of dedifferentiation rate *K*_*T*_. Note that a smaller NSCC cycling rate *K*_*N*_ makes the treatment more robust to dedifferentiation. **F** Confirmation of the result in a virtual cohort trial with 1000 patients. Decreasing both CSC and CSCC cycling rate leads to a higher fraction of cases with tumour shrinkage in the face of dedifferentiation.

To check the robustness of this synergy for different system parameters, we generate cohorts of 1000 virtual patients. For every patient, we sample a set of system parameters, and a value describing the strength of the applied treatment (see Section II). We then count the number of cases in which tumoural growth becomes negative if we only apply the treatment to the CSC compartment (Figures 3C blue line), to the NSCC compartment (orange line), and both compartments in parallel (green line). We repeat this for systematically altered dedifferentiation rates *K*_*T*_. In this way, we confirm that targeting both compartments in parallel is indeed less sensitive to dedifferentiation, even if system parameter values are randomly altered within biologically plausible ranges.

A similar synergy under standard parametrisation and in the virtual cohort can be observed in case of targeting both CSC and NSCC cycling rates *K*_*C*_, *K*_*N*_ (Figures 3D, 3E, 3F), however in the virtual cohort inhibiting cell cycling has turned out to be less successful than inducing apoptosis.

### C. Even under dedifferentiation, perturbing CSC-related parameters exerts the largest relative effect on tumour growth

In the previous section, we have shown that treatments which exert only one biological effect are more robust to dedifferentiation if they target both CSCs and NSCCs in parallel. Now, we want to compare the effectiveness of different treatments or treatment combinations. We start by comparing the effectiveness of single treatments.

The sensitivities under standard parametrisation for the case of all NSCCs dedifferentiating are plotted in Figure 4A for varying *K*_*T*_. In the absence of dedifferentiation, only CSC parameters (continuous lines) have a non-zero sensitivity (which may be easily confirmed analytically); however, increasing the dedifferentiation rate reduces the absolute magnitude of CSC parameter sensitivities and increases the absolute magnitude of NSCC parameters (dashed lines).

**FIG. 4:**
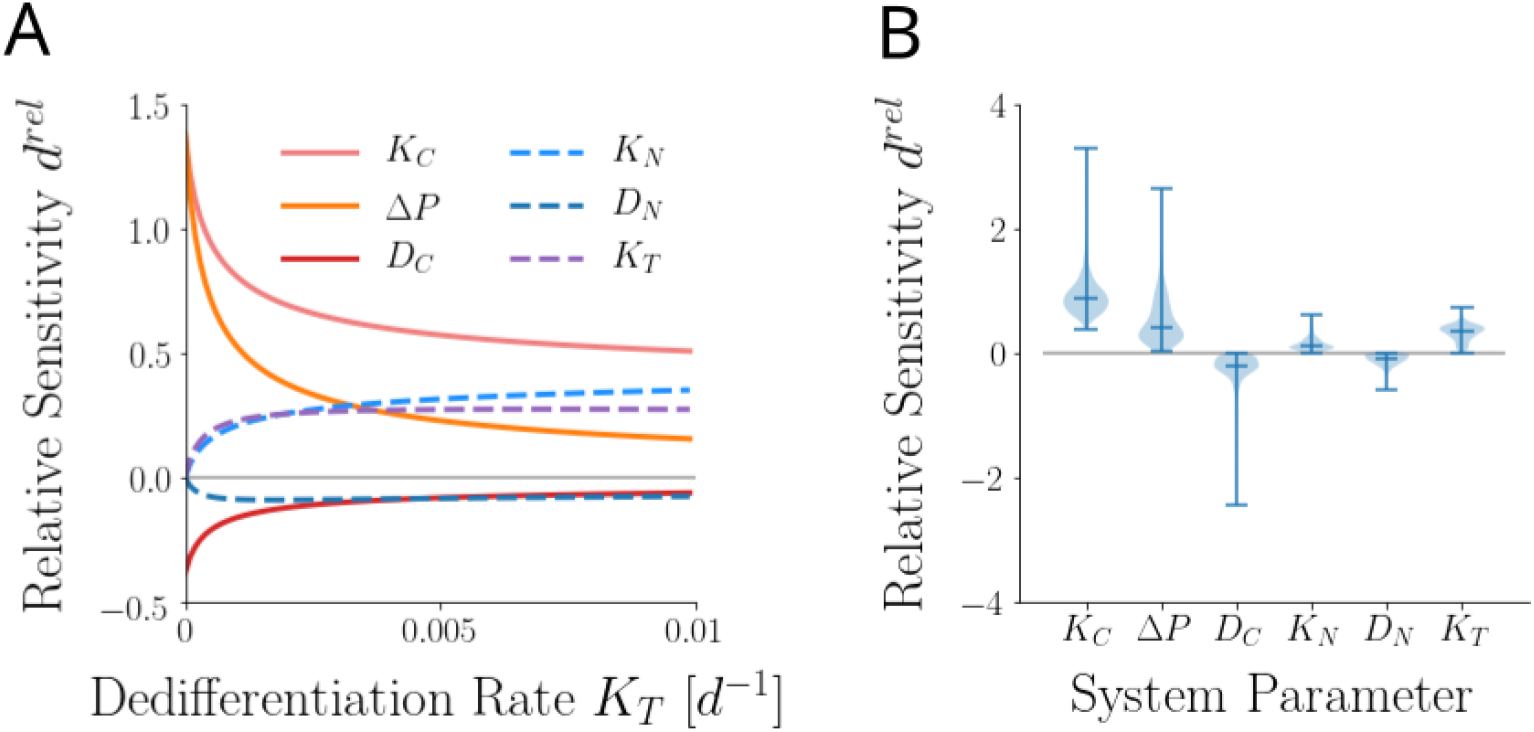
Sensitivity analysis of the system. **A** Numerical results under standard parametrisation for systematically varied dedifferentiation rates *K*_*T*_ : Dedifferentiation weakens the sensitivity of CSC-related parameters and strengthens the sensitivity of NSCC-related ones. **B** Distributions of system parameter sensitivities in a cohort of 1000 virtual patients with randomly drawn parameter values. Note that CSC-related parameters still are the most sensitive ones. In particular, CSC cycling rate *K*_*C*_ always shows a non-zero sensitivity.

To account for inter-patient variability, we again simulate a cohort of 1000 virtual patients as described before. The resulting distributions of sensitivities are plotted in Figure 4B. Interestingly, at the level of the complete cohort, CSC related parameters still exhibit larger absolute sensitivity coefficients than the NSCC related ones. In addition, the CSC cyling rate *K*_*C*_ is the only parameter that showed a distinctly non-zero sensitivity across all virtual patients.

### D. Treatments may interact in an antagonistic or synergistic fashion

Finally, we aim to quantify the strength of all pairwise interactions of two different treatments which affect different system parameters. In particular, we want to elucidate which pairs of treatments show a synergistic effect, where a combination treatment yields superadditive effects. Conversely, we also want to also identify all antagonistic treatment interactions, i.e. cases of a subadditive combined effect (see Figure 5A for an illustration).

**FIG. 5:**
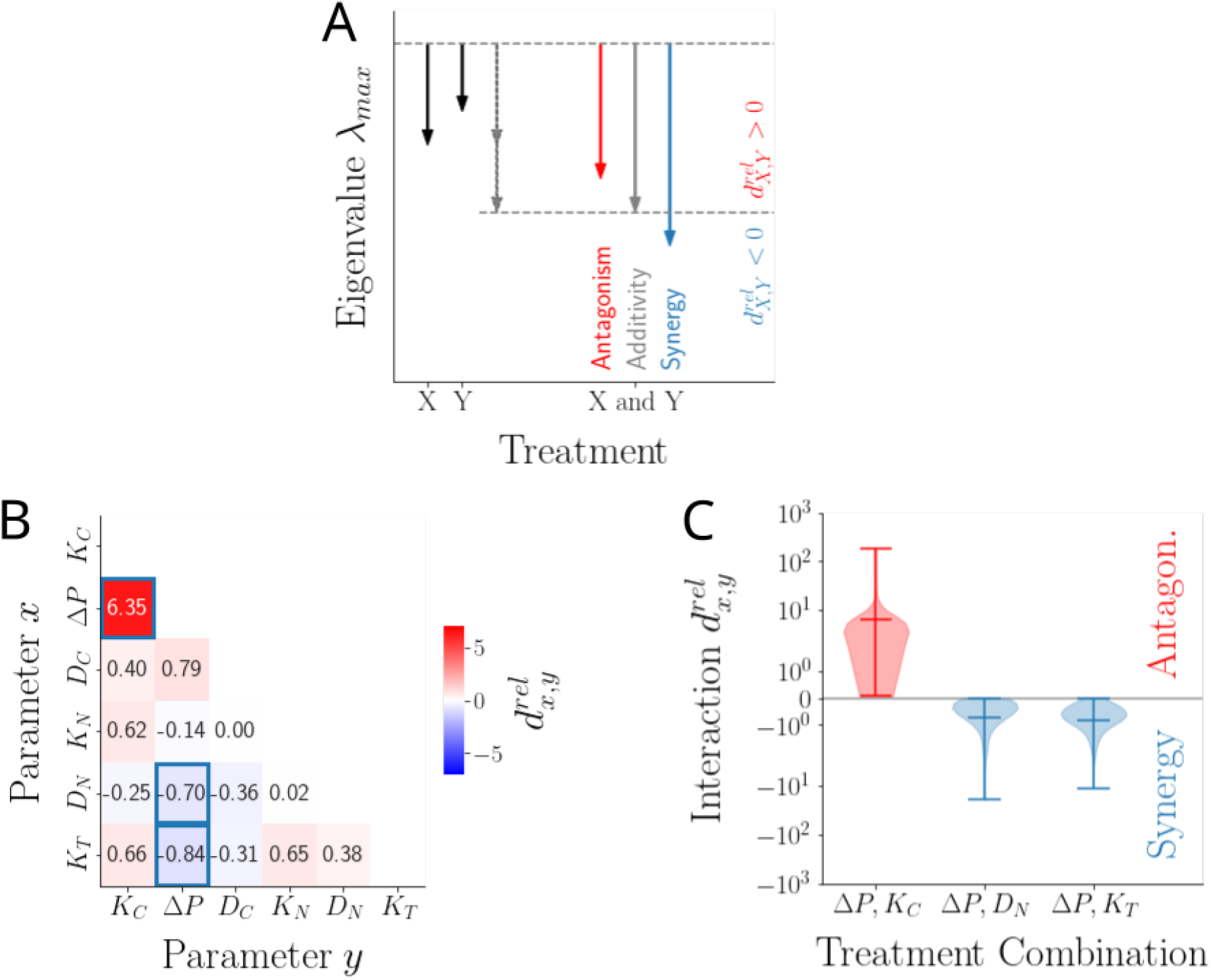
Treatment interactions in a cohort of 1000 virtual patients with randomly drawn parameter values. **A** Illustration of antagonism, additivity and synergy between two single treatments *X, Y*. **B** Mean of the interaction terms 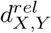for all treatment pairs *X, Y* across the cohort. Negative values (blue regions) denote synergy, i.e. superlinear effects; positive values (red regions) denote antagonism, i.e. sublinear effects. Three pairs of treatments are marked with cyan boxes. The distributions of their interaction values are plotted in panel C. **C** Distributions of 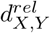for the three treatment pairs marked in panel B across the entire cohort. Note the consistent antagonistic interaction between CSC fate parameter Δ*P* and CSC cycling rate *K*_*C*_, as well as the consistent synergistic interaction between CSC fate parameter Δ*P* and NSCC apoptosis rate *D*_*N*_ and NSCC dedifferentiation rate *K*_*T*_, respectively.

If only NSCCs *N*_1_ dedifferentiate at rate *K*_*T*_ *>* 0, inspection of Equation 2 already suggests a complicated interaction structure between the system parameters. In particular, note that the nonlinearity of the root term couples stem cell with non-stem cell parameters in a way that does not lend itself to a simple analysis anymore. Allowing all NSCCs *N*_1_…*N*_*M*_ to dedifferentiate complicates these expressions even more. For this reason, we again numerically interrogate a cohort of virtual patients with random parameter values.

Throughout the cohort, we notice a strong antagonistic interaction between Δ*P* and *K*_*C*_, indicating that a combination of these two treatments will likely yield subadditive effects. (Indeed, this antagonism is straight-forward to show analytically for the case of *K*_*T*_ = 0 and thus not at all surprising, see Appendix D.) However, on average also some synergistic interactions become apparent, in particular between Δ*P* and *K*_*T*_, and between Δ*P* and *D*_*N*_. This suggests that targeting CSC fate Δ*P* in combination with either NSCC dedifferentiation rate *K*_*T*_ or apoptosis rate *D*_*N*_ will yield synergistic, superadditive reductions in tumoural growth rate (Figure 5B, cyan boxes). Plotting the complete distributions of these three interaction coefficients confirms that this is a consistent trend throughout the entire cohort (Figure 5C).

## IV. DISCUSSION

In this work, we extended a literature model [16] of the dynamics of a cancer stem cell (CSC) driven tumour to also account for cellular dedifferentiation. We revealed that dedifferentiation may contribute to carcinogenesis and that dedifferentiation increases the rate of tumoural growth. Both results are in accordance with pre-clinical, [23, 40] and clinical results ([41] and the references therein). Results from simpler, two-compartment toy models have confirmed these effects [26, 42]. Additionally, we revealed that older non-stem cancer cells (NSCCs) quantitatively contribute more strongly to these effects than younger ones. This suggests that not only suppressing cellular dedifferentiation, but also reducing NSCC life time and the number of NSCC cell divisions may be promising avenues for limiting tumoural growth *in vivo*.

The importance of NSCC behaviour has also been pointed out by Morton *et al*. [43] who used an agent-based cellular automaton model without dedifferentiation to show that the life span of NSCCs strongly modulates the progression of solid tumours. In contrast to our results, however, they showed that a longer NSCC lifespan may actually delay tumoural growth via spatial inhibition of CSC growth. In the same vein, Hillen *et al*. [44] used a partial differential equation model without dedifferentiation to show that removing NSCCs from a tumour can decrease competition for space and resources and thus increase overall tumour growth. Hence, *in vivo* dedifferentiating NSCCs might exert an additional, negative effect onto tumoural growth via competitive effects. Based on these results, Jilkine [45] has suggested that a successful treatment needs to selectively target CSCs over NSCCs, because otherwise tumoural growth speed and survival probability may increase. However, our results demonstrate that under dedifferentiation NSCCs stimulate tumoural growth and thus targeting them may actually prove to be beneficial. Experimental studies (see below) agree and suggest that these positive effects indeed seem to predominate.

Next, we showed that our model replicates the clinical observation that treatment-resistant CSCs cause long-term tumour survival despite an initial reduction in tumour size [5, 46]. Thus, we turned to the question of specifically targeting CSCs. Importantly, we showed that dedifferentiation enables a tumour to evade treatments directed at CSCs, because treatments need to exert significantly stronger effects to successfully offset even small amounts of dedifferentiation. We demonstrated that this becomes especially severe if a treatment solely targets the CSC compartment without affecting NSCCs. Conversely, we revealed that a treatment exerting the same biological effect in both CSCs and NSCCs at the same time is more robust towards dedifferentiation, and thus more likely to be successful. We further confirmed this synergy by showing its presence in a numerically simulated cohort of virtual patients with randomly drawn parameters from biologically plausible ranges. Taken together, this reveals an important caveat of the paradigm of exclusively directing therapy at eradicating CSCs [6] and suggests that treatment strategies directed at both CSCs and NCSCs may be more effective in the face of dedifferentiation.

This result can be expected to be even more relevant in the light of the fact that the tumour microenvironment can stimulate NSCC dedifferentiation [47], for instance in situations of cellular stress such as hypoxia [48]. Importantly, results from experiments and mathematical modelling suggest that in melanoma a reduction of CSCs below a certain threshold stimulates NSCC dedifferentiation [49], which, as argued by Jilkine [45], further casts doubt on the success of therapeutic strategies based on CSC eradication alone.

Finally, we aimed to prioritise among different treatment avenues by calculating the sensitivities of all individual model parameters. We observed that dedifferentiation decreases the sensitivity of CSC-related parameters and increases the sensitivity of NSCC-related ones. Nonetheless, CSC-related parameters generally remained the most sensitive parameters, highlighting their continued importance for tumoural treatment even in the face of cellular dedifferentiation. In addition, we quantified all pairwise treatment interactions throughout our virtual cohort. We demonstrated the existence of antagonistic, as well as synergistic treatment interactions; in particular, we showed that targeting CSC cycling rate and CSC fate might lead to a sublinear treatment outcome. In contrast, a treatment targeting CSC fate seems to interact synergistically with a treatment that increases NSCC apoptosis rate or decreases NSCC dedifferentiation rate. This further reinforces the idea of synergistic effects in targeting CSCs and NSCCs in unison as described above.

Research on cancer cell lines and xenograft mouse models has already pointed into the same direction. For the case of breast cancer, Hirsch *et al*. [50] have demonstrated that a specific combination of metformin, which selectively targets CSCs, and doxorubicin, a chemotherapeutic agent that only kills NSCCs, reduces tumour mass and prolongs remission much more effectively than either drug alone. Similarly, in prostrate cancer, selectively targeting CSCs with the de-methylating agent 5-azathioprine or the signalling inhibitor *γ*-tocotrienol in combination with selectively targeting NSCCs with an enhancer of androgen receptor degradation leads to significant synergy in tumour suppression and therapeutic efficacy [51]. Another promising trend in currently ongoing preclinical research lies in identifying suitable single substances which target both CSCs and NSCCs in parallel [52–54].

Altogether, our study demonstrates from a theoretical point of view the various effects of cellular dedifferentiation on the growth and treatment avenues of tumours, highlighting the biological importance of this process. Our results have revealed a number of insights which we hope to be helpful in guiding future molecular and clinical research. In particular, we anticipate that future studies on using inhibitors of dedifferentiation, on limiting the proliferative potential of NSCCs, and on combining CSC- and NSCC-focused treatments will prove to be useful in the clinic. More broadly, our results stress the importance of understanding the behaviour of non-stem cancer cells next to the cancer cell compartment in a growing tumour, both from a basic research point of view as well as from an applied perspective.

## ACKNOWLEDGMENTS

M.M.F. acknowledges funding by the Deutsche Forschungsgemeinschaft (DFG, RTG2424 CompCancer).

## AUTHOR DECLARATIONS

### B. Code availability

All source code for the reproduction of the numerical analyses is available in the following github repository: https://github.com/Matthias-M-Fischer/Tumourgrowth

### C. Author contributions

Both authors conceived of the presented ideas; M.M.F. carried out model derivations and analyses, and produced the initial version of the manuscript, both with help from N.B. N.B. contributed to the final version of the manuscript and supervised the project. All authors have read and approve of the final version of this manuscript.

## Appendix A: Approximation of the leading eigenvalue of the system

First, observe that System 1 can be written as

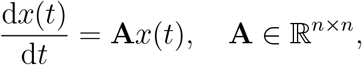

where **A** = **A**_**0**_ + **Ã** with

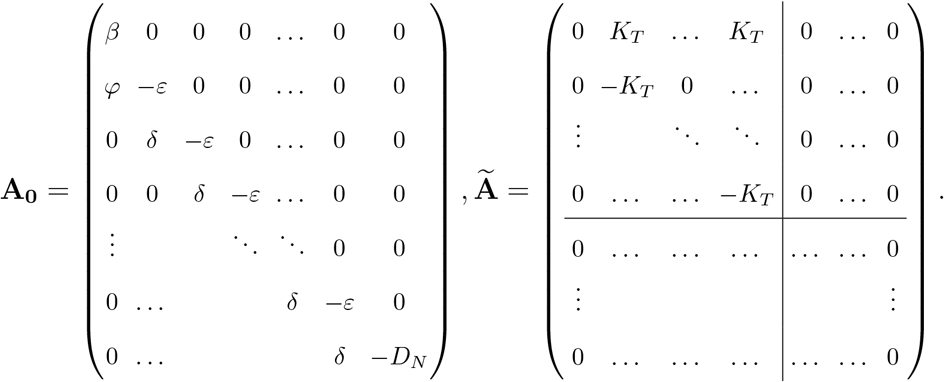

The lower-triangular matrix **A**_**0**_ represents the system without dedifferentiation, and **Ã** contains the changes that arise due to dedifferentiation. Note, that dedifferentiation only happens up to a certain arbitrary, but fixed NSCC cell compartment *N*_*d*_, after which dedifferentiation stops.

Clearly, **A**_**0**_ has a simple eigenvalue *λ*_1_ = *β* with left eigenvector *u* = [1, 0, 0, …, 0] and the right eigenvector

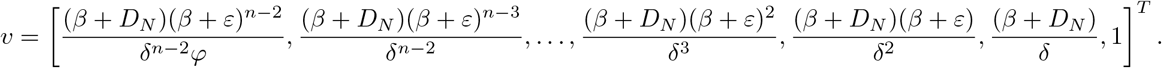

Let 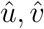denote the normalised eigenvectors. Classical analytical matrix perturbation theory (see for instance Theorem 4.4 from [55], for a proof refer to page 149f therein), allows us to express the change of *λ*_1_ which arises as a consequence of this perturbation, as

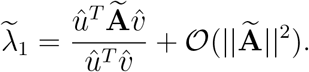

Neglecting higher order terms, this reduces to

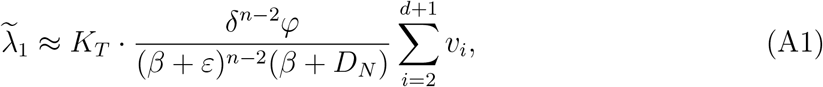

where *v*_*i*_ represents the *i*-th element of the (non-normalised) vector *v*.

Since 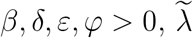 is a positive and increasing linear function of *K*_*T*_.

The prefectors *φ/*(*β* + *D*_*N*_) and the sum term over *v*_*i*_ are constants determined by the system parameters. By assuming that the realised *in vivo* growth rate *β* of the CSC compartment and the NSCC apoptosis rate *D*_*N*_ are small compared to the NSCC cycling rate *K*_*N*_ (see Section II), we find for the increase 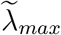in the lading eigenvalue *λ*_*max*_ that

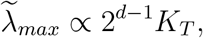

indicating a sharp linear increase of *λ*_*max*_ with a slope that is exponential in the number of NSCC divisions.

## Appendix B: Targeting only CSCs can quickly fail in the face of dedifferentiation

### 1. The case of *d* = 1

We begin our examination by considering the model in which only *N*_1_ cells can dedifferentiate.

Assume first that we want to target *D*_*C*_. Let *ω* = *D*_*C*_ *− K*_*C*_Δ. In the absence of dedifferentiation, any *ω >* 0 will lead to treatment success. However, if *K*_*T*_ *>* 0, we require that

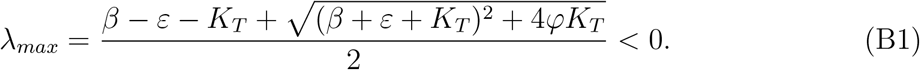

Expressing *D*_*C*_ in terms of *K*_*C*_Δ*P* and *ω* in *β* of Equation B1 yields

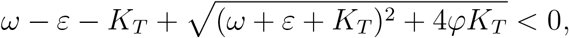

which reduces to

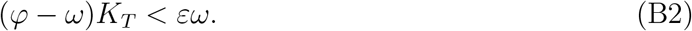

This always holds if *ω ≥ φ*, which is equivalent to *D*_*C*_ *> K*_*C*_. However, if *ω < φ*, then treatment will succeed if and only if

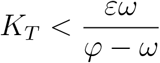

Next, assume we want to target *K*_*C*_. Let *ω* = *K*_*C*_Δ*P/D*_*C*_. Without dedifferentiation, any *ω <* 1 will lead to treatment success. For *K*_*T*_ *>* 0, however, we express *K*_*C*_ in terms of *ω* and the other two CSC parameters, and plug this expression into *β* and *φ* of Equation B1.

In this way, we find that a sufficiently strong treatment of *ω ≤* Δ*P* will succeed regardless of *K*_*T*_, however a weaker treatment (Δ*P < ω <* 1) will only be successful as long as

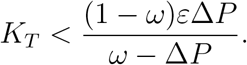

Finally, if we want to target Δ*P*, we let *ω* = Δ*PK*_*C*_*/D*_*C*_. Without dedifferentiation, the treatment succeeds for any *ω <* 1. If, however, *K*_*T*_ *>* 0, we express Δ*P* in terms of *ω* and the other two CSC parameters. Plugging this into B1 yields:

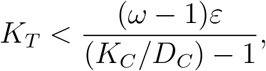

where *D*_*C*_ *< K*_*C*_. Thus, a sufficiently strong treatment of *ω <* 1 will always succeed, but for any weaker treatment there exists a critical positive value for *K*_*T*_, where the treatment begins to fail.

### 2. The case of arbitrary *d*

A similar set of arguments can also be made for the case of arbitrary *d*, i.e. the case where multiple NSCCs can dedifferentiate. For this, we use the first-order approximation of the eigenvalue given by Equation A1. Importantly, the approximation only holds for sufficiently small values of *K*_*T*_, and only as long as *β* remains the leading eigenvalue of **A**_**0**_, i.e. as long as *β > −D*_*N*_. Slightly rearranging for clarity and considering only the last summand of the sum over the eigenvector elements, we get the lower bound

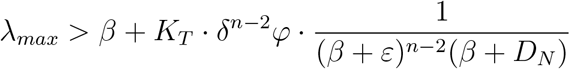

Applying any treatment directed at the CSC compartment will reduce *β*, however *δ*^*n−*2^*φ* will stay constant and since *β > −D*_*N*_ the fraction will stay positive. Thus, a sufficiently large dedifferentiation rate *K*_*T*_ can cause *λ*_*max*_ to exceed zero, and thus the treatment to fail.

If *−D*_*N*_ *− K*_*N*_ *< β < −D*_*N*_, the leading eigenvalue of **A**_**0**_ will switch to *−D*_*N*_. By calculating its associated left and right eigenvalue, we can derive a similar approximation as before, again showing that a sufficient increase in *K*_*T*_ may render a treatment directed at the CSC compartment unsuccessful by causing *λ*_*max*_ to exceed zero. For the sake of brevity, we here omit the details.

Finally, if *β < −D*_*N*_ *−K*_*N*_, the leading eigenvalue of the system will become *−D*_*N*_ *−K*_*N*_ with an algebraic multiplicity of *n −* 2. Thus, for *n >* 3, the eigenvalue is not unique anymore, and thus the approximation cannot be calculated.

## Appendix C: Targeting CSCs and NSCCs in parallel is more robust to dedifferentiation

Here, we examine whether targeting both CSCs and NSCCs in parallel can be more effective.

### 1. The case of *d* = 1

We start with the model in which only *N*_1_ cells can dedifferentiate (*d* = 1). Consider again Equation B2, describing the necessary and sufficient condition for treatment success if the CSC apoptosis rate *D*_*C*_ is target of the treatment. Expanding all aggregate parameters and separating the CSC and NSCC parameters yields

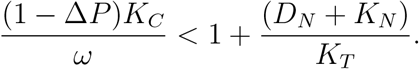

Clearly, additionally increasing NSCC apoptosis rate *D*_*N*_ expands the region in parameter space in which treatment is successful; thus, targeting both CSCs and NSCCs can be expected to prove beneficial. Interestingly, the same holds for increasing NSCC cycling rate *K*_*N*_, which might seem counterintuitive at first glance. However, this finding does make sense, because in the examined model only *N*_1_ cells dedifferentiate. Thus, increasing *K*_*N*_ leads to NSCCs leaving this compartment faster, and thus reduces cellular dedifferentiation.

Similar arguments for synergy can be made for the cases where not *D*_*C*_, but Δ*P* or *K*_*C*_ is targeted using the same approach based on considering the necessary and sufficient condition for the success of the respective treatment. Because these calculations are entirely analogous to the previous one, we omit them here.

### 2. The case of arbitrary *d*

For the case of an arbitrary number *d* of dedifferentiating compartments, we can use the Gershgorin circle theorem [39] on the complete coefficient matrix **A** in a column-wise manner to derive bounds on all system eigenvalues. We here only consider the case of *d* = *M*, i.e. all NSCC compartments dedifferentiate, since all other cases 1 *< d < M* follow completely analogously.

From the first column, we get the region *S*_1_ = [*β − φ, β* + *φ*]. Columns 2 … *n −* 1 yield the region *S*_2_ = [*δ − ε −* 2*K*_*T*_, *δ − ε*]. Finally, the last column yields *S*_3_ = [*−D*_*N*_ *−* 2*K*_*T*_, *−D*_*N*_]. Since *∀i* : *λ*_*i*_ *∈ 𝒰*_*j*_*S*_*j*_, we get that *∀i* : *λ*_*i*_ *<* max*{β* + *φ, δ − ε, −D*_*N*_ *}*. Expanding aggregate parameters, we thus get

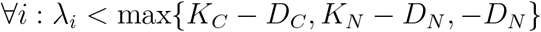

Assume a CSC-treatment which is sufficiently strong, so that *K*_*C*_ *− D*_*C*_ *<* 0. Then it is still not guaranteed that the treatment succeeds in the face of dedifferentiation as long as *K*_*N*_ *− D*_*N*_ *>* 0. (In fact, numerical simulations presented in the main part of this manuscript show that treatment may fail very easily even for small values of the dedifferentiation rate *K*_*T*_.) However, additionally applying a NSCC-treatment which ensures that *K*_*N*_ *− D*_*N*_ *<* 0 will always ensure treatment success regardless of dedifferentiation rate *K*_*T*_.

## Appendix D: Antagonism between Δ*P* and *K*_*C*_ for *K*_*T*_ = 0

In the absence of dedifferentiation (*K*_*T*_ = 0), we get by inspection of the leading eigenvalue that only *K*_*C*_ and Δ*P* can be expected to interact. A reduction of *K*_*C*_ to *η*_1_*K*_*C*_ causes an absolute reduction of the leading eigenvalue by Δ_1_ = Δ*PK*_*C*_(1 *− η*_1_), and similarly a reduction of Δ*P* to *η*_2_Δ*P* causes an reduction of Δ_2_ = Δ*PK*_*C*_(1 *− η*_2_), where *η*_1_, *η*_2_ *∈* (0; 1). Applying both parameter alterations in unison causes an absolute eigenvalue reduction of Δ_12_ = Δ*PK*_*C*_(1 *− η*_1_)(1 *− η*_2_).

By direct calculation, it is easy to show that

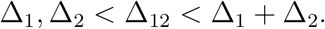

In other words, the two treatments will always interact antagonistically, and their combined effect will be strictly smaller than the sum of the two single effects., however larger than the effect of either of the two treatments alone.

